# Why clinical trials are terminated

**DOI:** 10.1101/021543

**Authors:** Theodore R. Pak, Maria D. Rodriguez, Frederick P. Roth

## Abstract

**Background:** Evidence-based clinical practice relies on unbiased reporting of negative results. Meta-analysis of drug safety and efficacy across many clinical trials is difficult given the unconstrained nature of reasons that are provided to ClinicalTrials.gov to explain clinical trial terminations.

**Methods and Findings:** We scanned all trials in ClinicalTrials.gov marked with the “terminated” status (N=3122), meaning the trial had been stopped before the scheduled end date. Under the current reporting framework, any number of reasons may be given for termination, and these need not conform to a controlled vocabulary. Here we develop a controlled vocabulary for trial termination, and map each terminated trial to as many as three vocabulary terms. Mapping to this “ontology of termination” allows further analysis and conclusions. First, we identify the subset of terminated trials that ended citing safety concerns (6.2%) or failure to establish efficacy (10.8%), and were further able to stratify these rates across trials of different phases. Second, we examine termination reasons where a stricter data model could have preserved more evidentiary value, either because the data model was misused (7.6%) or because the given reason left unclear whether the decision to terminate was based on analysis of the data (74.9%, with 20.4% mentioning a decision-maker that may have had access to the data). Third, we show that imposing a controlled vocabulary of reasons for termination would avoid ambiguity and improve the evidentiary value of clinical trials.

**Conclusions:** We encourage wider use of an “ontology of termination” and propose four questions that should be posed on trial termination. These simple steps would promote transparency and enable ready access to negative trial results for meta-analysis.

## Introduction

Evidence-based practice has established itself as the appropriate method of incorporating scientific research results into the practice of medicine. However, evidence-based practice relies on the medical literature accurately reflecting the current set of evidence for and against any given scientific theory—particularly, the efficacy of some proposed treatment against an indication. The various players in the industrial-academic network of pharmaceutical producers, clinical researchers, and publishers have interests that do not always align, and often have motives that are contrary to the ideal of openness underlying modern science. When this impacts results presented to other researchers and clinicians, evidence-based practice as a whole is compromised. Selective reporting creates biases that not only threaten current clinical practice and undermine guideline recommendations, but can cultivate further biases that derail future research [1].

One phenomenon that influences selective reporting is now known as “positive-outcome bias,” which is the higher probability that studies showing favorable or statistically significant outcomes will be publicized in medical journals and conferences. Positive-outcome bias has been demonstrated in randomized trials of selection of article drafts for publication by peer review [2] and selection of abstracts for presentation at a scientific meeting [3] (collectively termed “publication bias”). Likewise, researchers are less likely to publish negative outcomes [4–6], which can be considered “outcome-reporting bias.” There also may be incentives to suppress adverse events that arise during a clinical trial of a treatment, which preclude building an accurate profile of harms—a profile that is critical for making appropriate treatment choices [7]. At its worst, distortion of harms has been deliberately orchestrated to commercialize drugs at the expense of patient safety [8, 9]. While not always this explicit, a review of publications of randomized clinical trials has found substantial variability in the reporting of harm-related results [10].

Public identification of all clinical trials and their protocols has been advocated as a way of alleviating public concern over undisclosed safety problems with drugs [11]. The United States enacted legislation to mandate registration of all trials testing effectiveness of investigational drugs for “serious or life-threatening” conditions with the Food and Drug Administration (FDA) Modernization Act, section 113 (FDAMA 113) in 1997 [12]. In 2000, the National Library of Medicine on behalf of the National Institutes of Health implemented this registry with the website ClinicalTrials.gov [13, 14]. In 2004, the International Committee of Medical Journal Editors (ICMJE) announced that any clinical trial must be registered by September 2005 in a public clinical trials registry such as ClinicalTrials.gov as a prerequisite for publication in any of its journals. The number of trials registered within ClinicalTrials.gov increased by 73% in the following five months [15]. Because most clinical trials with a U.S. testing site must now be registered by law and as a typical prerequisite for publication, ClinicalTrials.gov has become useful for cross-disciplinary analysis of trends in clinical trial protocol and conduct [16–19]. Some of these studies have found areas where ClinicalTrials.gov’s protocols could be improved to facilitate further transparency [16].

The transparency offered by a mandated database such as ClinicalTrials.gov offers the possibility of a source of representative negative as well as positive results. Negative results are not only useful for fair meta-analysis of treatment options, but can also illuminate which compounds have been proven safe in humans by Phase 1 trials but are simply awaiting research to show them to be effective for an indication [20]. The strong arguments that positive results are over-represented in published clinical literature [1, 21] are reflected in surveys of the publication characteristics of trials registered in ClinicalTrials.gov; while 78.3% of publications related to registered clinical trials claimed positive outcomes [16], surveys of the drug development pipeline show that 39%-64% of drugs actually advance to the next step of each phase of clinical trials [22]. Only recently in September 2007, a new mandate was enacted by the FDA Amendments Act (FDAAA) for registrants to update ClinicalTrials.gov with full trial results within one year of the trial completion [23, 24]. Given the small number of trials for which this mandate currently applies, we sought to identify “golden negatives” in the rest of the database.

ClinicalTrials.gov allows for a trial to persist in one of several states, of which the “Terminated” designation within the 

~~~
overall_status
~~~

 field indicates that a trial was halted before its scheduled endpoint. In combination with the 

~~~
why_stopped
~~~

 field, where registrants can provide a reason for the termination, negative results can be inferred. We sought to examine the suitability of these two fields in the ClinicalTrials.gov database for finding negative trial results. The 

~~~
why_stopped
~~~

 field is freeform, which means that clarity and evidentiary utility of the reasons provided varied considerably. While parsing the set of reasons, we developed an ontology that may be useful for classifying clinical trial terminations. Implementing and requiring use of this (or a further simplified) ontology in a mandatory registration database such as ClinicalTrials.gov would provide a high-quality repository of negative evidence and thus benefit rigorous meta-analysis of results and evidence-based clinical practice.

## Methods

90,523 records of clinical trials were downloaded on May 31, 2010 from ClinicalTrials.gov as “full study descriptions” in XML format. All studies with a 

~~~
why_stopped
~~~

 field (N=3122) were separated and filtered into a database and randomly ordered. Figure 1 shows the incidence of notable words in the corpus of 

~~~
why_stopped
~~~

 text as a word cloud, with often-observed sequences such as “interim analysis” combined into phrases, obvious spelling errors corrected, and English stop words and lesser-interesting phrases removed (e.g., “the,” “due to,” “lack of “).

**Figure 1:**
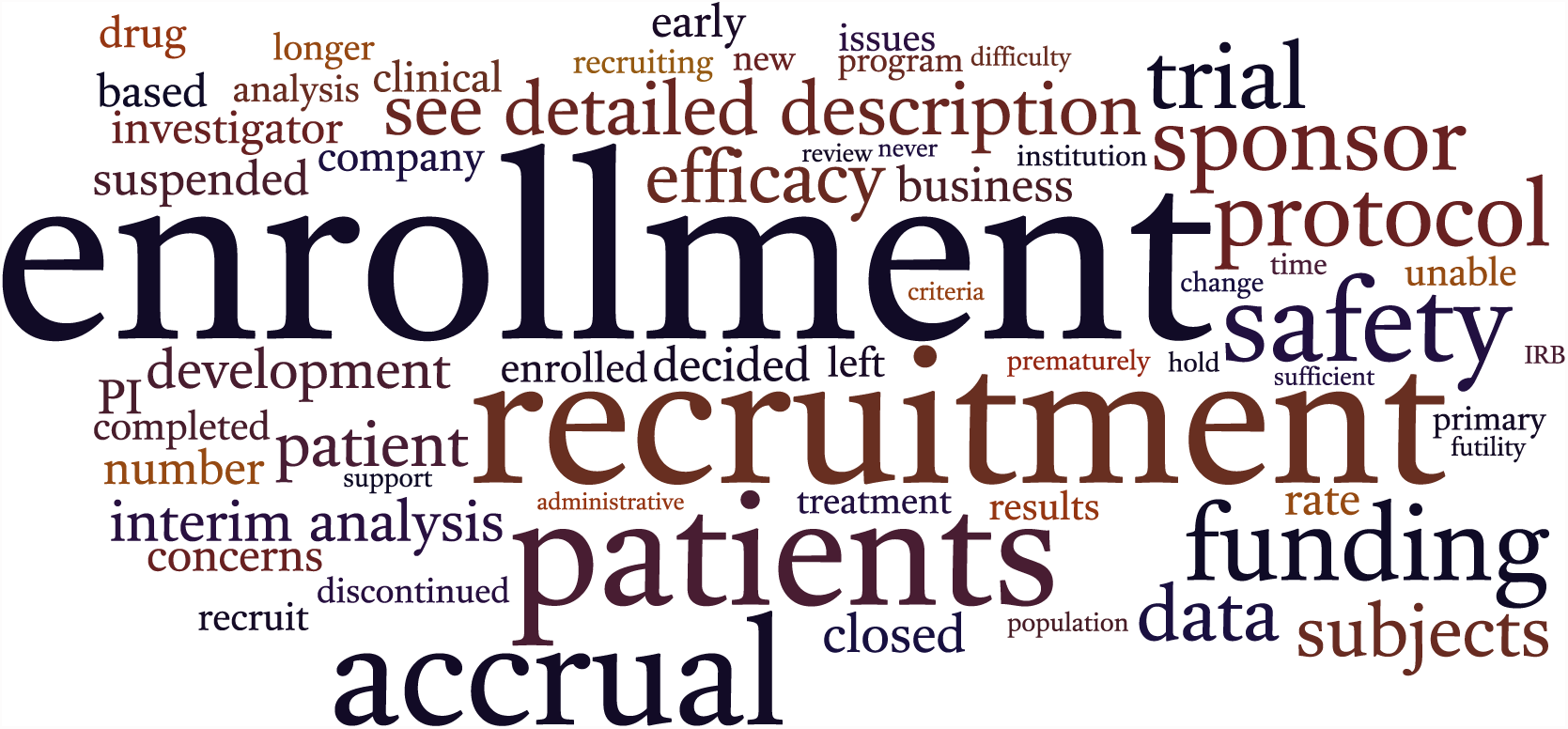
Word cloud generated from why_stopped text. The incidence of the 60 most frequent notable words and phrases are shown. Font size is proportional to the frequency of the word or phrase, and position and color are arbitrary. Often-observed sequences such as “interim analysis” were combined into phrases, obvious spelling errors were corrected, and English stop words and lesser-interesting phrases were removed (e.g., “the,” “due to,” “lack of “).

Patterns in the frequency of words—for instance, the prominence of “enrollment” and logically related words like “recruitment” and “patients”—suggest that a classification system for termination reasons can be constructed. Accordingly, each 

~~~
why_stopped
~~~

 value was labeled by researchers with up to three terms from an ontology that was iteratively assembled with the goal of defining a small vocabulary that could capture the essence of the vast majority of the free-form reasons. This controlled vocabulary is displayed in Table 1.

**Table 1:**
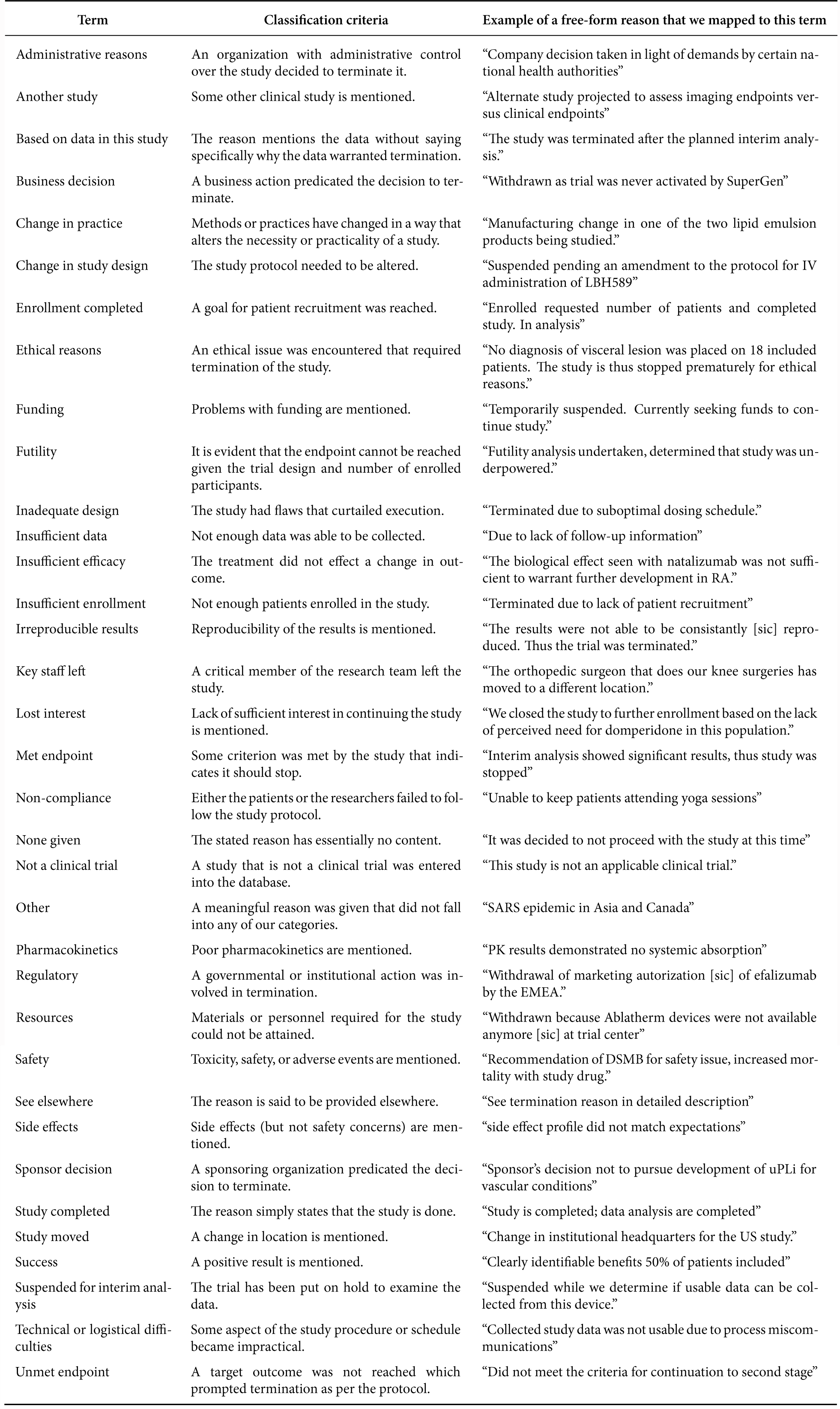
The “ontology of termination” created and used for this study.

Each given reason was also assessed according to four additional criteria: *i*) did researchers indicate that the data had been examined? *ii*) can we infer the trial had never started? *iii*) was efficacy ruled out as a factor in the termination? and, *iv*) was safety explicitly ruled out as a factor in the termination?

We remained blind to the title, sponsor, purpose, and description of the trial while labeling reasons with ontology terms. To ensure accuracy and consensus in the application of each term, each term assignment for each given reason was reviewed by one or two of us. Conflicts in labeling were discussed until consensus was reached.

For each termination, we also extracted from the trial database the 

~~~
phase
~~~

 field and the 

~~~
nct_id
~~~

 field allowing each termination to be linked back to its original record, if necessary. The 

~~~
phase
~~~

 field allows a trial to span multiple phases, such as “Phase 3/Phase 4”; we conservatively rounded these down to the lowest provided phase number for all ensuing analysis. If no 

~~~
phase
~~~

 was specified, we classified it as “Other.”

Some of the 

~~~
why_stopped
~~~

 values stated that a reason was given elsewhere, typically in the 

~~~
de-tailed_description
~~~

 field. For these trials, we first applied the ontology term “See elsewhere” and then manually searched ClinicalTrials.gov for the reason using the study’s 

~~~
nct_id
~~~

. If the reason was not found, we searched the history of the trial record to find the archived version that contained the reason, and noted the date when it had been edited out of the record.

Next we distilled our applied ontology into six categories that represent the evidentiary value represented by each term, with respect to the result of the trial being negative, positive, or among various degrees of uncertainty about the trial outcome. Each category was assigned a priority so that reasons assigned to multiple terms from different categories could be resolved into one category. We determined this priority by weighing the relative informational strength of each category; for example, a “negative efficacy” reason in tandem with a “neutral” reason conveys an overall negative finding, so the “negative efficacy” category takes priority. The categories, priorities, and their respective terms are displayed in Table 2.

**Table 2:** Categories for each term in our ontology. The priority assigned to each category was used to map reasons with terms from multiple categories to one category; the category with priority 1 takes the greatest precedence, the category with priority 2 takes the second greatest precedence, and so on.

| Category | Priority | Terms |
| --- | --- | --- |
| Negative efficacy | 1 | Based on data in this study, Futility, Insufficient efficacy, Pharmacokinetics, Unmet endpoint |
| Positive | 2 | Success |
| Negative safety | 3 | Safety, Side effects |
| Possibly negative | 4 | Administrative reasons, Business decision, Funding, Irreproducible results, Lost interest, Sponsor decision |
| Neutral | 5 | Another study, Change in practice, Change in study design, Ethical reasons, Inadequate design, Insufficient data, Insufficient enrollment, Key staff left, Met endpoint, Non-compliance, Other, Regulatory, Resources, See elsewhere, Study moved, Suspended for interim analysis, Technical or logistical difficulties |
| Misuse | 6 | Enrollment completed, None given, Not a clinical trial, Study completed |

Some of the categories in Table 2 deserve special mention. We decided to distinguish between reasons that most likely provide “neutral” evidentiary value from those that were “possibly negative.” Reasons that specified a direct external circumstance that led to trial termination were classified as “neutral,” while reasons leaving open the possibility that termination was based (at least in part) on insufficient efficacy or safety were classified as “possibly negative.” Our criteria for qualifying a nonnegative, non-positive, and safety-unrelated term as “possibly negative” as opposed to “neutral” was if the term implicated the decision-making of a party that may have had access to the data generated in the course of the clinical trial, e.g, a sponsor, industry collaborator, or administrative body.

The “misuse” category was attached to terms that do not constitute valid reasons to terminate according to the protocols of ClinicalTrials.gov. A registrant should not be marking their trial with the 

~~~
overall_status
~~~

 “Terminated” if they have merely completed enrollment or if the study is completed. In the former case, the “Active, not recruiting” status is more appropriate, and in the latter case, the “Completed” status is more appropriate [25]. The “None given” term indicates that the 

~~~
why_stopped
~~~

 value was entirely non-informative, e.g., it simply restated that the study was terminated without providing a reason. The “Not a clinical trial” term indicates that a registrant had created a record in ClinicalTrials.gov for a study that was not an applicable clinical trial.

The software package R version 2.11.0 was used for all statistical analyses. Wordle (http://www.wordle.net/) was used to generate Figure 1 and the Protovis library (http://www.protovis.org/) was used to generate Figure 9.

## Results

As shown in Figure 2, most reasons were only labeled by one term from our ontology. This is not surprising due to the typically short length of the reasons; ClinicalTrials limits the length of the 

~~~
why_stopped
~~~

 field to 160 characters [25], and we observed a mean of 7.8 words (standard deviation ±5.7) per reason.

**Figure 2:**
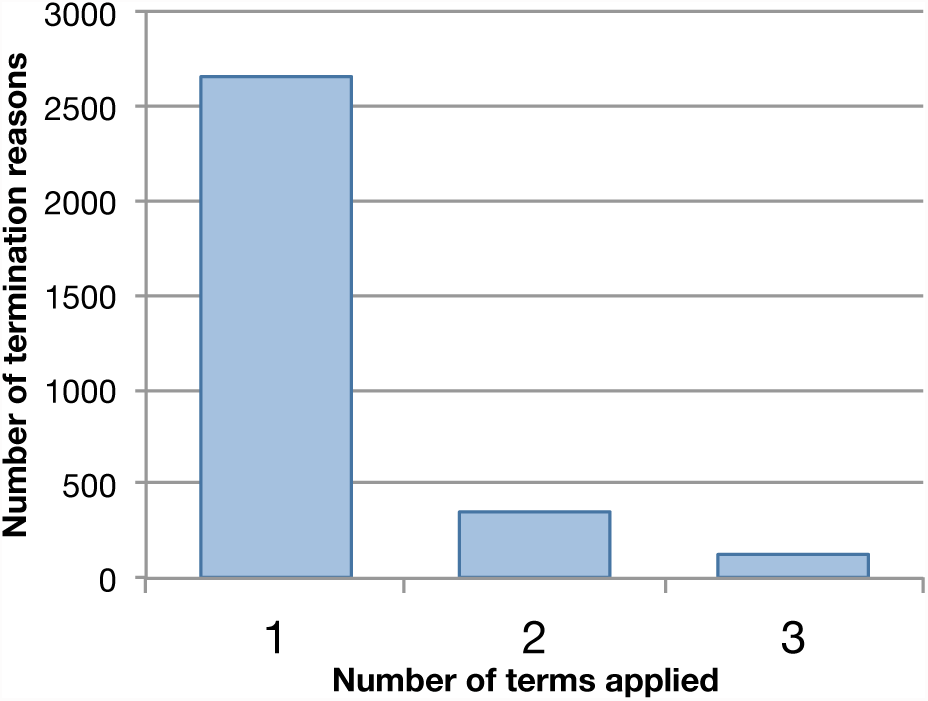
Distribution of ontology terms mapped to termination reasons. A total of 3623 assignments of a controlled vocabulary were made, to a total of 3122 given reasons.

Figure 3 shows the distribution of term counts for termination reasons from different phases of clinical trials. Most notable is the greater likelihood of a Phase 3 termination reason receiving multiple labels, indicating that Phase 3 reasons were usually more complicated or more thoughtfully described than reasons for other phases.

**Figure 3:**
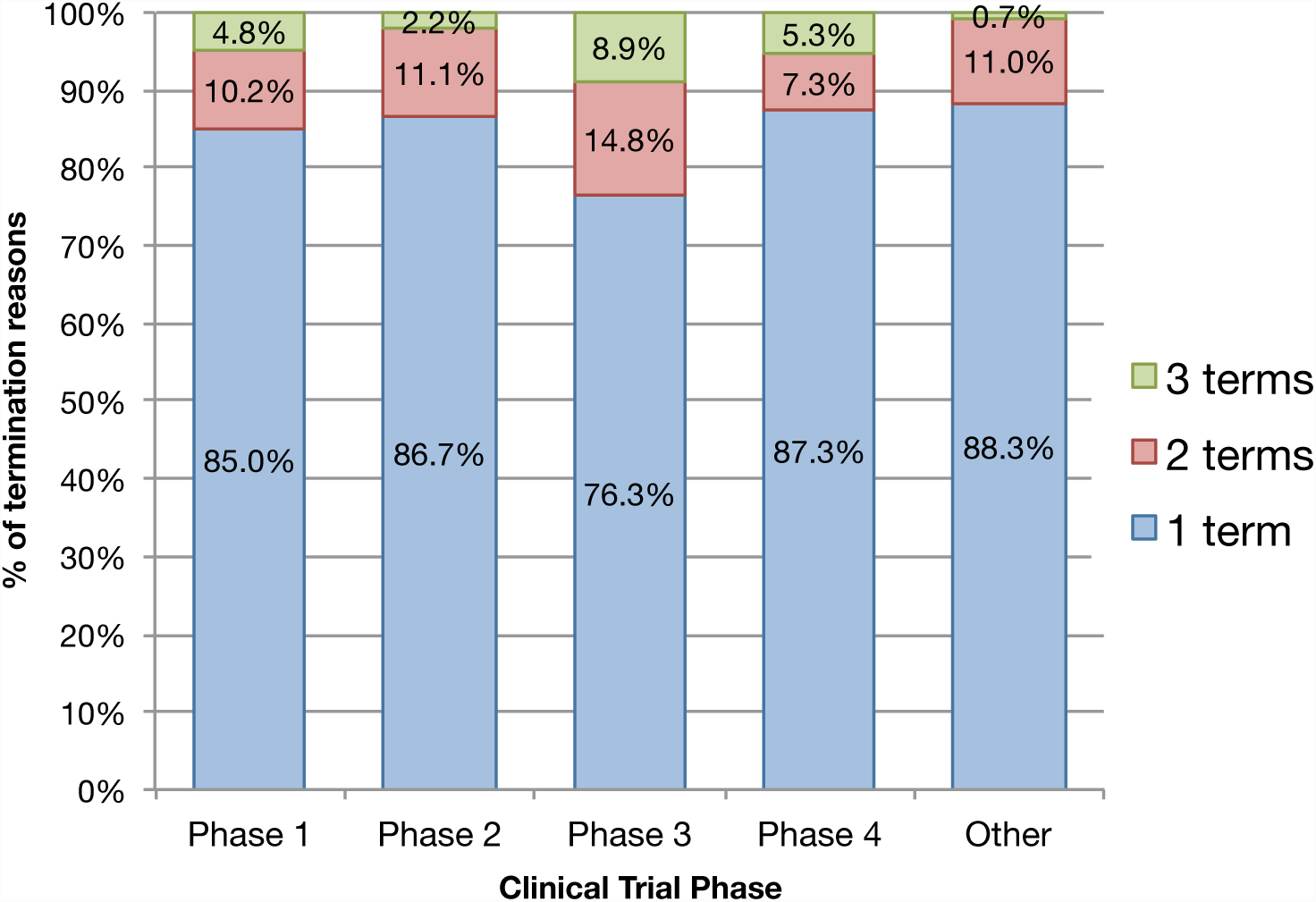
Percentage of terminations at each phase labeled with either one, two, or three ontology terms. Total counts are: Phase 1, N=479; Phase 2, N=881; Phase 3, N=575; Phase 4, N=449; and N=738 for no specified phase.

Figure 4 shows the frequency of each term within our dataset. By far the most common term applied was “insufficient enrollment”, mentioned in 33.7% of termination reasons and consequently categorizing a wide swath of terminations as “neutral”. The most common “possibly negative” reasons were “funding” and “business decision”, appearing in 7.6% and 7.3% of reasons, respectively. “Insufficient efficacy” was the most common “negative efficacy” reason, appearing in 6.8% of reasons; “Safety” appeared in a nearly equivalent 6.7% of reasons. A sizable 5.0% of reasons contained no usable information at all. All other terms appeared in less than 5.0% of responses.

**Figure 4:**
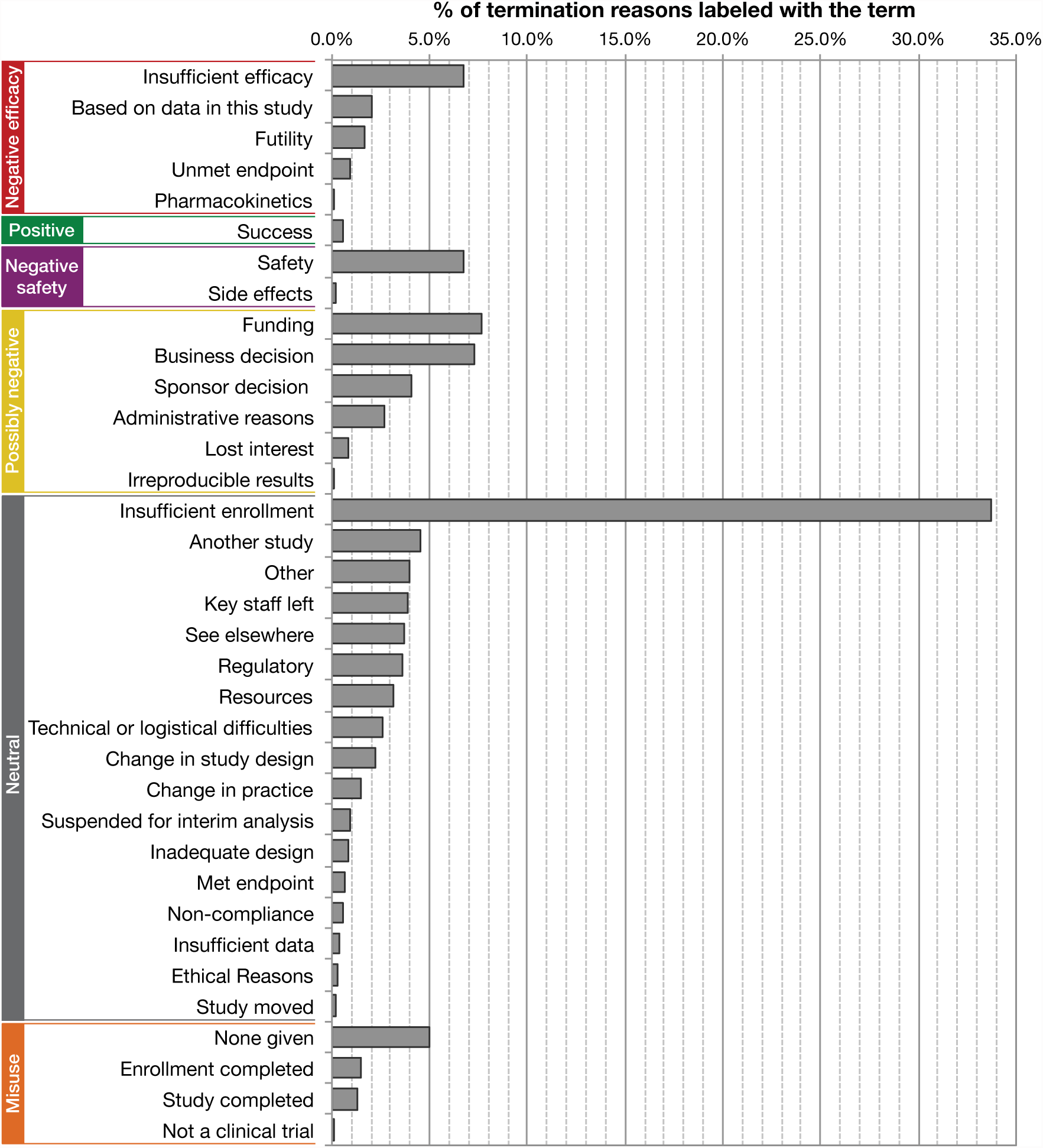
Percentage of termination reasons labeled with each ontology term. Terms are sorted by category and then by percent incidence. (N=3122)

While applying ontology terms to reasons, we also evaluated each reason according to four criteria assessing whether interim data analysis may have precipitated the termination (see **Methods**). A summary of assignments according to these criteria appears in Figure 5. As before, Phase 3 trials are remarkable for displaying the most extreme qualities: they are the most likely to terminate asser the researchers have looked at the data, least likely to terminate before starting, and most likely to explicitly state that termination was not due to efficacy or safety concerns.

**Figure 5:**
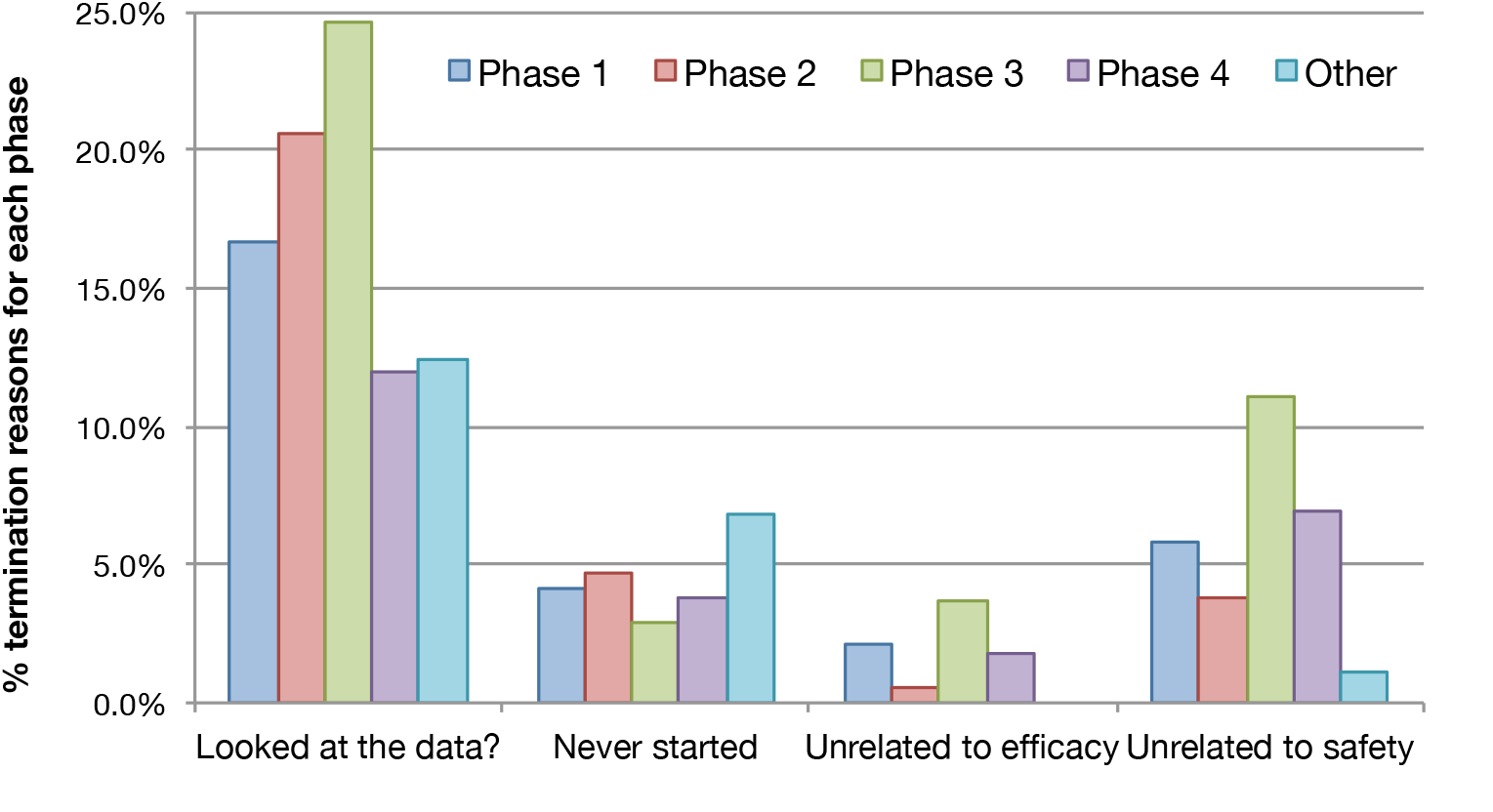
Percentage of terminations at each phase flagged for four criteria. The four criteria are described in more detail in **Methods**. Total counts for each phase are the same as in Figure 3.

Using the categories and priorities shown in Table 2, we classified each termination reason given according to its evidentiary value, summarized in Figure 6. Positive reasons were the most scarce at only 0.5%. About 10.8% or 337 trials could be classified as “golden negatives” that terminated due to failure to establish efficacy, while 6.2% of trials reported termination primarily due to safety concerns.

**Figure 6:**
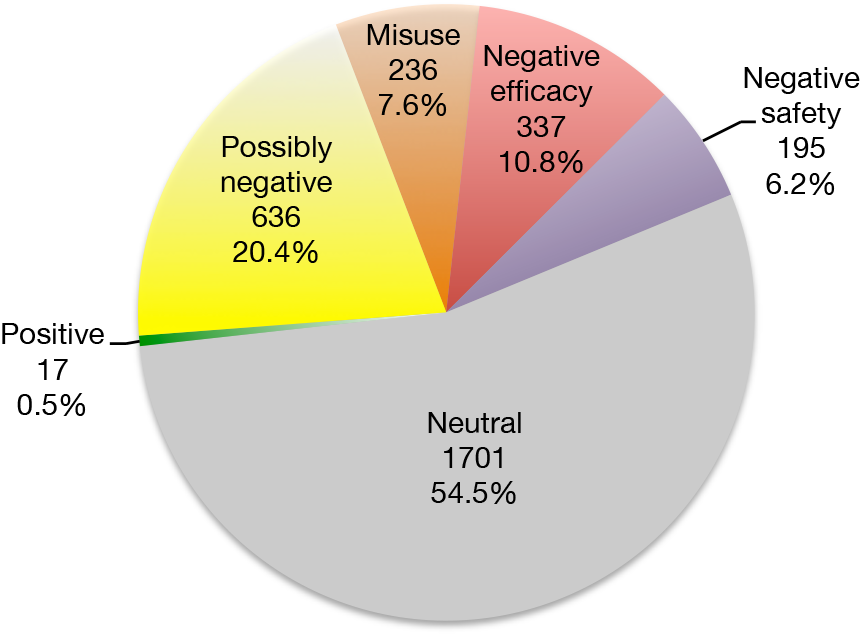
Number and percentage of terminations that fell into six categories of evidentiary value. These categories are defined in Table 2. (N=3122)

For the remaining 82.5% of terminated trials, the reasons provided did not clearly communicate whether termination was based on analysis of the clinical trial data. “Neutral” reasons predominated, comprising 54.5% of terminations. 20.4% of terminations were deemed “possibly negative,” because the decision to terminate involved a party that may have had access to trial data. A substantial fraction (7.6%) of reasons were only assigned terms that we consider to be misuses of the 

~~~
why_stopped
~~~

 field.

Separating the categorized reasons for each phase of clinical trials produces the chart shown in Figure 7. Phase 3 displays the greatest incidence of termination for negative reasons (19.5%), the least incidence of termination for neutral reasons (44.5%), and also the greatest incidence of termination for positive reasons (1.6%). Phase 1, 2, and 3 trials had similar incidences of “possibly negative” termination reasons (22.3%, 20.2%, and 22.6%, respectively). From Phase 1 through Phase 3, an upward trend can be observed in the incidence of termination for both negative and positive reasons, while a downward trend can be observed in the incidence of termination for neutral reasons and negative safety reasons.

**Figure 7:**
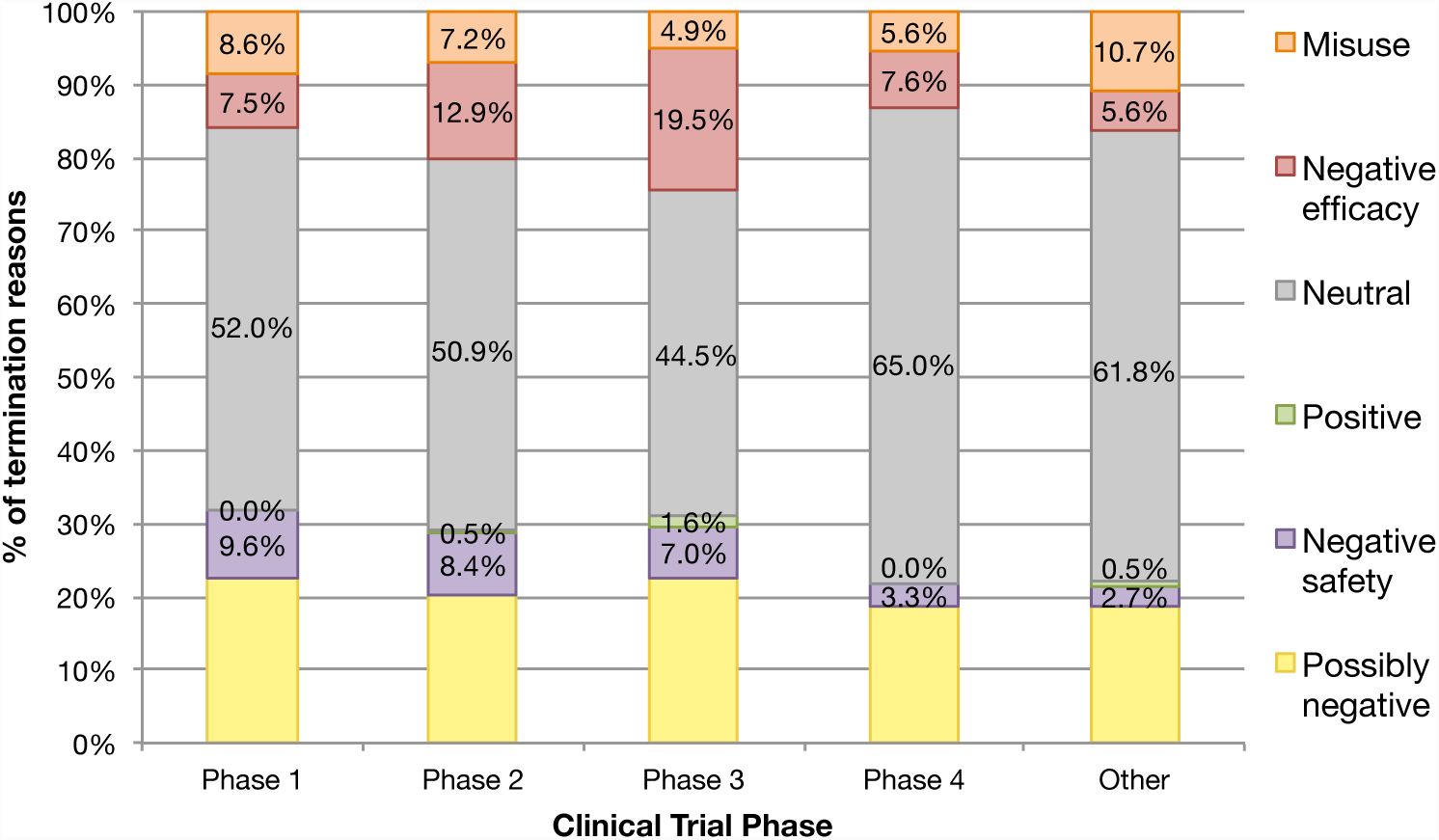
Percentage of terminations at each phase that fell into six categories of evidentiary value. These categories are defined in Table 2. Total counts for each phase are the same as in Figure 3.

Figure 8 revisits the four criteria examined in Figure 5, examining their correlation with each of the categories of evidentiary value. As might be expected, a very high proportion of “positive” and “negative efficacy” terminations gave reasons indicating that the investigators had examined the data, with about half of “negative safety” terminations doing the same. “Neutral” reasons were rarely flagged by any of the criteria. “Possibly negative” reasons rarely indicated that the data was examined, and also rarely said explicitly that safety and efficacy were not factors in termination. “Negative efficacy” reasons were most likely to include the assertion that safety was not involved in terminating the trial, followed closely by “possibly negative” reasons; this perhaps indicates that trial coordinators, when unable to show efficacy but not finding unexpected safety issues, wish to allay concerns about safety where further trials are contemplated.

**Figure 8:**
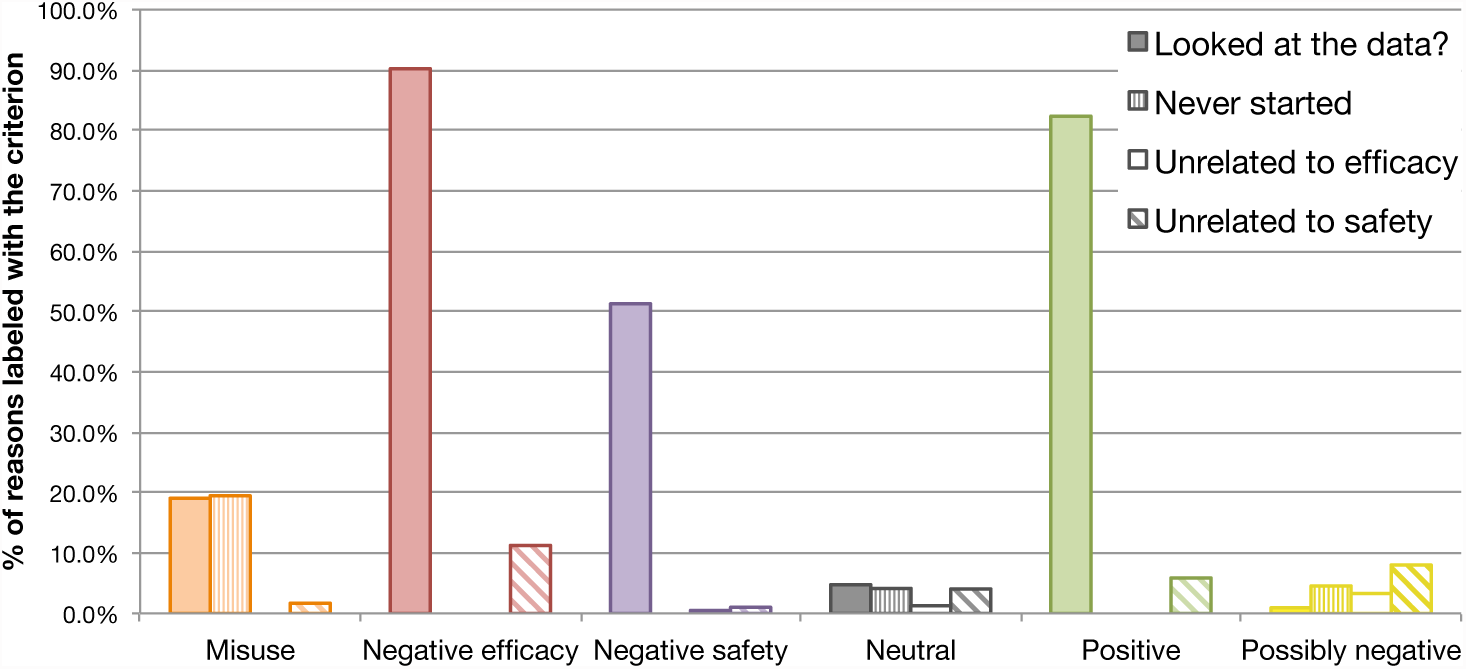
Percentage of terminations in each category of evidentiary value flagged for four criteria. The criteria are described in more detail in Methods. Total counts for each category are displayed in Figure 6.

While most reasons were linked to only one ontology term (Figure 2), we examined reasons labeled with multiple terms to explore the correlations between causes for termination. Figure 9 represents these correlations as a node-link graph where term frequency is expressed as node size and correlation strength as the width of the lines between each node. Correlation strength was calculated using the Jaccard index—the ratio of the number of terminated trials labeled with both terms to the number of terminated trials labeled with at least one of the terms. Only correlations significant at *p* < 0.05 according to Fisher’s Exact Test are shown. Twelve terms did not correlate significantly with any other term and were dropped from the diagram, leaving one well-connected giant component. “Insufficient enrollment,” the most commonly applied term, exhibited significant connections to nineteen other terms.

**Figure 9:**
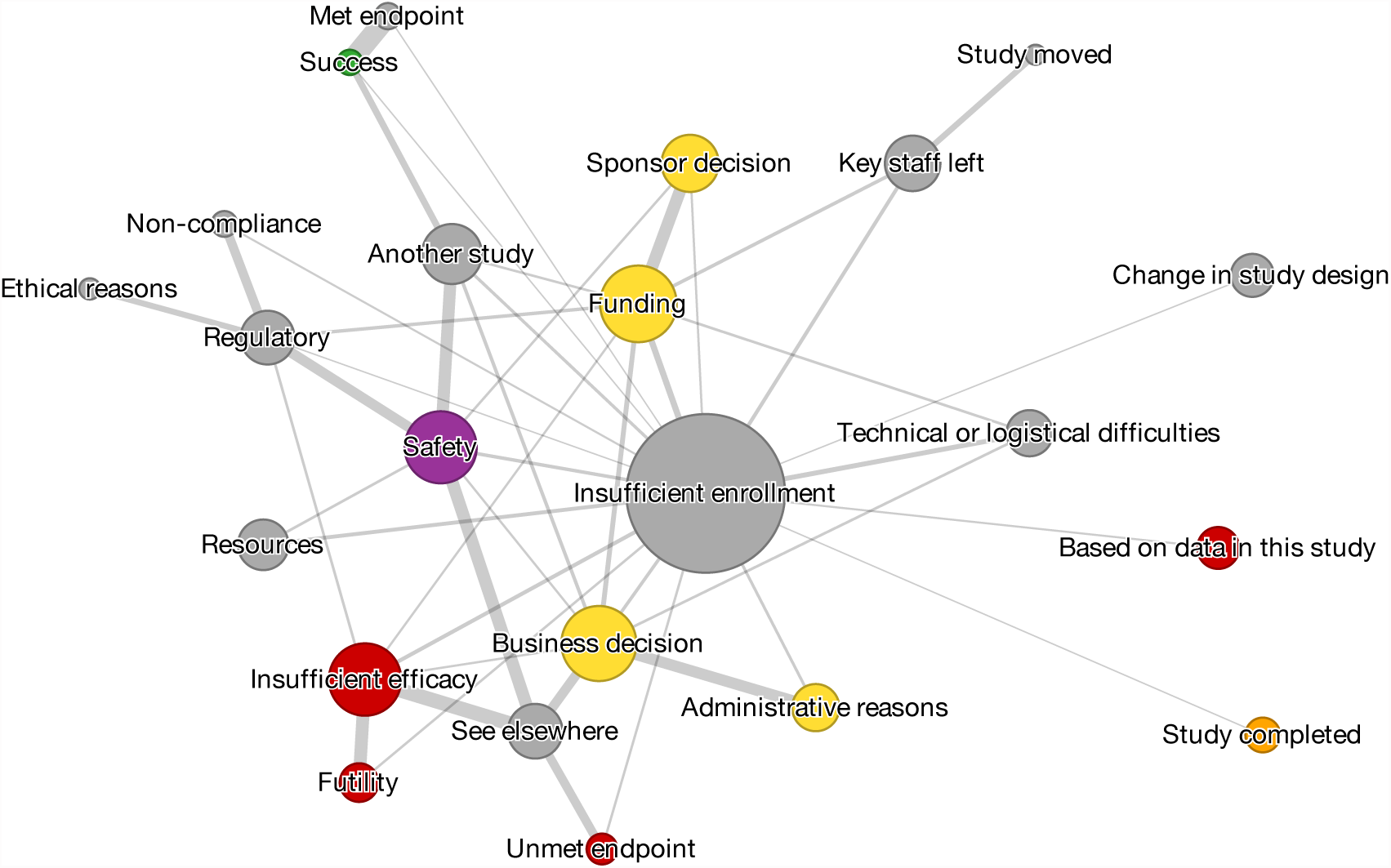
A node-link graph showing significant correlations between assigned terms in the “ontology of termination.” Nodes are grouped by color using the categories in Table 2 and are sized according to overall term incidence (see Figure 4). The width of the links between each node reflects the correlation strength between the two terms. Only correlations that are significant at the *p* < 0.05 level according to Fisher’s exact test are shown. 12 ontology terms have no significant correlations and were removed from the diagram.

A strong correlation is observed between “futility” and “insufficient efficacy.” This correlation is sensible—interim trial results can point to a reduced upper bound of efficacy. A reduced upper bound on efficacy in turn leads to an increase in the estimate of the number of additional subjects that would be required to provide sufficient statistical power to distinguish such an efficacy value from that of the null hypothesis. Where the reduced estimate of efficacy suggests that not enough subjects are available to establish significance, it may be considered futile to continue the trial. The yellow nodes, all from the “possibly negative” category, cluster with each other and “neutral” terms more strongly than with either “negative safety” or “negative efficacy” terms. Interestingly, the “see elsewhere” term correlates strongly with “negative efficacy,” “negative safety,” and “possibly negative” terms, perhaps reflecting either registrants’ reluctance to enter such reasons into the 

~~~
why_stopped
~~~

 field or their belief that the field is too constrained to adequately capture the complexity of their reason.

## Discussion

### Primary findings

We labeled all terminated clinical trials in the ClinicalTrials.gov database with terms from an “ontology of termination,” producing subsets of trials that carried either positive, negative efficacy, negative safety, possibly negative, or neutral evidence for myriad combinations of treatments and indications. Such a trial categorization permits systematic analysis. For example, not only can a reviewer assess a fuller history of evidence regarding a drug’s inefficacy or safety issues, but drugs that have failed at any particular phase can be extracted for renewed experimentation against other relevant indications [20]. This idea of “repurposing” drugs, which has become increasingly attractive as thousands of old drug patents lapse and the average cost of approving new compounds rises into the billions of dollars [26, 27], could be facilitated by computational models that predict drug efficacy for arbitrary indications. Such models would require unbiased training sets derived from past experimental evidence.

Surprisingly few terminations (10.8%; Figure 6) clearly indicated negative efficacy findings; however, this still provides a set of 337 trials with negative outcomes for efficacy and 195 trials (6.2%) with negative safety outcomes for future evaluation and meta-analyses (Tables S1 and S2, supplementary data). For 82.5% of terminated trials, the reasons provided did not allow a determination of whether the trial was terminated in whole or part based on data. We do not wish to fault the registrants for this, except perhaps in the 7.6% of cases where the entries are frank misuses of the data model. Although we argue here that provided reasons should contain evidentiary value where possible, ClinicalTrials.gov registrants were not specifically required to maximize this potential value when populating the 

~~~
why_stopped
~~~

 field.

Trends in clinical trial termination separated for clinical trials in different phases can be observed. These correlate well with the nature of those phases. Phase 3 trials are extraordinarily expensive, at a mean cost of $86 million, more than three times the average cost of Phase 2 trials, which are in turn often more expensive than Phase 1 trials [27]. The stakes are also higher for Phase 3 trials, in the sense that two successful Phase 3 trials can represent the final hurdle for treatment approval by the FDA [20]. Therefore, they are less likely to be terminated without good and carefully considered reasons (Figure 3). These trials are also least likely to terminate before starting (Figure 5), which is to be expected given that more effort, planning, and resources are dedicated toward their completion. By Phase 3 the drug has presumably demonstrated both safety and efficacy so trial registrants are more likely to state that these are not reasons for termination (Figure 5). Figure 5 also shows the increasing willingness of investigators to look at the data before terminating trials in phases 1 through 3. Figure 7 shows trends for more “negative efficacy” reasons for termination, more “positive” terminations, less “misuse” terminations, and less “neutral” terminations as trials progress from phases 1 through 3, correlating the increased cost and risk of each phase with a greater likelihood of continuing the study until a result is obtained. Since every phase involves evaluation of safety factors, it is also expected that terminations for “negative safety” reasons decreases steadily from phases 1 through 4 (Figure 7). Phase 4 does not follow many of the other trends of the phases. Because these trials take place after the drug is FDA-approved, seeking data on the drug’s optimal use, there are different motivations and incentives for outcome and completion.

Most terminations appear to be the result of factors not involving interim analysis of the results of the trial and therefore do not immediately implicate bias. Enrollment is the primary problem (Figure 4), but there are myriad other contributing reasons given, such as precedence by other studies, logistical and resource issues, inappropriate trial design, resources or staff being unavailable, and regulatory intervention. Even for these trials, however, the reasons given generally do not allow us to determine whether interim analysis of trial data was performed. Disillusionment with interim data could, for example, decrease the motivation for resource allocation and patient recruitment. Overall, these “neutral” reasons are responsible for about half of registered trial terminations (Figure 6).

A surprisingly large number of reasons (20.4%, or 636 trials total, Figure 6) implicated decision-making by parties likely to have access to the trial’s interim data without being specific about why that party is no longer willing to support the trial. Occasionally, additional details were added that added insight into the third party’s considerations, e.g., “patent legal settlement” or “marketing reasons”, but these details cannot exclude the possibility that the third party based their decision in whole or in part on the interim trial data, and such details were provided too infrequently to be catalogued in the ontology. The only way we could divine from the given reason whether the trial data itself was a factor in the decision would be an explicit statement by the registrant that either *i*) the trial was never started so that data could not have been a factor, or *ii*) efficacy or safety did not play a role in the decision to terminate. As shown in Figure 8, very few registrants giving “possibly negative” reasons noted specifically that the termination was not based on trial data.

Finally, 236 registrants (7.6% of all terminations, Figure 6) either failed to provide any reason at all, or misused the “Terminated” designation for the 

~~~
overall_status
~~~

 field. Such entries dilute the value of the ClinicalTrials.gov database and call for more stringent validation of entries before they become part of the official record. In particular, it should not be possible for 5.0% of terminations to lack a substantive reason. A response such as “Trial cancelled” completely defeats the purpose and evidentiary value of the 

~~~
why_stopped
~~~

 field. Secondly, misusing the “Terminated” status when a study has closed enrollment or is complete may seem like benign error, but it could be used to conceal a negative outcome or skirt the mandatory reporting of results enacted by the FDAAA [23]. Consequently, the possibility of misuse of the status field should also be eliminated.

### Recommendations for ClinicalTrials.gov

In addition to more stringent regulation of the “Terminated” status and the 

~~~
why_stopped
~~~

 field, the high rate of “possibly negative” reasons we uncovered indicates that transparency and clarity would only be achieved if ClinicalTrials.gov supplemented or replaced the free-form 

~~~
why_stopped
~~~

 field.

We would suggest, at a minimum, providing investigators with explicit questions as to whether *i*) the trial was started, *ii*) the data was either directly or indirectly examined, *iii*) an interim evaluation of efficacy was a factor in the termination, and *iv*) whether an interim evaluation of safety was a factor in termination. Such queries, if truthfully answered, would effectively sort all “possibly negative” reasons into either the “neutral”, “negative efficacy”, or “negative safety” categories. Consistent and mandated reporting will improve the evidentiary value of registered trial terminations.

For even more detail, a field where the registrant selects all applicable terms from our proposed “ontology of termination” would clarify the ambiguities currently present in the free-text format. Otherwise, we predict that meta-analysis of clinical trial terminations will remain labor-intensive and inaccessible to most consumers of ClinicalTrials.gov. Compounding this problem is the current allowance for registrants to specify that readers should “see elsewhere”, usually in 

~~~
de-tailed_description
~~~

, for the reason—an option that is unfortunately explicitly recommended by the NIH’s instructions for registrants [25]. Not only does this confuse and complicate consumption of the database, as the 

~~~
detailed_description
~~~

 field is another larger free-text field primarily intended for other data, but 

~~~
detailed_description
~~~

 is already designated to contain the overflow of no less than five other fields [25]. Furthermore, it leaves the information more prone to being later removed from the 

~~~
detailed_description
~~~

 field without alerting administrators of the registry, as we discovered to have occurred in 9 of the 112 “see elsewhere” reasons that we investigated. In these cases, the history of the trial record had to be plumbed for the archived version that contained information on termination, and the free-text nature of this field makes automation of this process impracticable. Although 8 of these 9 subsequently-deleted termination reasons turned out to be “insufficient efficacy,” we do not assert that registrants are currently using this to intentionally obscure negative results; however, the potential for abuse certainly exists. Therefore, we would discourage ClinicalTrials.gov from promoting or allowing overloading of any of their data fields, and to revise the data model so that this is never required of a registrant.

### Strengths and limitations of this study

We assumed good faith on the part of all registrants when reviewing ClinicalTrials.gov data, so whatever a termination reason stated or implied was taken at face value. Our primary purpose in this study was to distill further evidentiary value from clinical trial terminations, and as a result this may have colored our ontology and the subsequent categorization priorities associated with it. We believe that this objective is sufficiently desired by researchers and the proponents of public clinical trial registries to serve as a fundamental design principle for the ontology. However, studies focused on other aspects of clinical trial termination may desire an ontology that is more or less weighted toward other aspects of the spectrum of reasoning. We also recognize that the reasons for termination can be complex, combining estimates of safety and efficacy with marketing, intellectual property and competition issues. Nonetheless, we believe it is reasonable to ask whether information about efficacy and safety that were gleaned from the trial were contributing factors.

### Conclusions

The premature termination of a clinical trial is an event that should be closely tracked by clinical trial registries. Termination can indicate a range of outcomes from extreme positive or negative results to innocent logistical difficulties, so registries should ensure that an informative reason for termination is provided. Without governments and journal-editors mandating registration, investigators would have little incentive to disclose the often uninformative or negative results of trial terminations. Fortunately, ClinicalTrials.gov already contains a large repository of data on terminated trials that can be analyzed to produce statistics on why and how clinical trials fail, and rescue the evidentiary value of true negative results for meta-analysis. To our knowledge this study is the first to describe an “ontology of termination” and its application to a clinical trial registry. Our hope is that this ontology will inform future analyses of clinical trial termination and encourage improvements in the transparency offered by trial registries by enumerating all possible factors that should be assessed when recording trial termination reasons. Increased transparency and clarity in clinical trial reporting should improve meta-analyses of clinical trial data, which serves to boost the efficacy of evidence-based practice.

## Acknowledgements

The authors wish to thank Jason M. Johnson of Merck Pharmaceuticals for the suggestion to pursue meta-analyses of the ClinicalTrials.gov data, and for insightful comments on the manuscript. We also thank Rahul C. Deo and other members of the Roth Laboratory for their valuable comments.

### Financial disclosure

FPR was supported by NIH grants (HG003224, HG004756, CA130266, MH087394 and HG004233), a Canadian Institute for Advanced Research Fellowship and a Canada Excellence Research Chair. No funding bodies had any role in study design, data collection and analysis, decision to publish, or preparation of the manuscript.

### Author contributions

FPR and TRP conceived and designed the study. FPR, MDR and TRP designed the “ontology of termination” and performed mapping of termination reasons to ontology terms. TRP analyzed the results. TRP drafted the manuscript with revisions by FPR.

### Competing interests

The authors declare no competing interests.

